# Structure of M^pro^ from COVID-19 virus and discovery of its inhibitors

**DOI:** 10.1101/2020.02.26.964882

**Authors:** Zhenming Jin, Xiaoyu Du, Yechun Xu, Yongqiang Deng, Meiqin Liu, Yao Zhao, Bing Zhang, Xiaofeng Li, Leike Zhang, Chao Peng, Yinkai Duan, Jing Yu, Lin Wang, Kailin Yang, Fengjiang Liu, Rendi Jiang, Xinglou Yang, Tian You, Xiaoce Liu, Xiuna Yang, Fang Bai, Hong Liu, Xiang Liu, Luke W. Guddat, Wenqing Xu, Gengfu Xiao, Chengfeng Qin, Zhengli Shi, Hualiang Jiang, Zihe Rao, Haitao Yang

## Abstract

A new coronavirus (CoV) identified as COVID-19 virus is the etiological agent responsible for the 2019-2020 viral pneumonia outbreak that commenced in Wuhan^1–4^. Currently there is no targeted therapeutics and effective treatment options remain very limited. In order to rapidly discover lead compounds for clinical use, we initiated a program of combined structure-assisted drug design, virtual drug screening and high-throughput screening to identify new drug leads that target the COVID-19 virus main protease (M^pro^). M^pro^ is a key CoV enzyme, which plays a pivotal role in mediating viral replication and transcription, making it an attractive drug target for this virus^5,6^. Here, we identified a mechanism-based inhibitor, N3, by computer-aided drug design and subsequently determined the crystal structure of COVID-19 virus M^pro^ in complex with this compound. Next, through a combination of structure-based virtual and high-throughput screening, we assayed over 10,000 compounds including approved drugs, drug candidates in clinical trials, and other pharmacologically active compounds as inhibitors of M^pro^. Six of these inhibit M^pro^ with IC_50_ values ranging from 0.67 to 21.4 μM. Ebselen also exhibited promising antiviral activity in cell-based assays. Our results demonstrate the efficacy of this screening strategy, which can lead to the rapid discovery of drug leads with clinical potential in response to new infectious diseases where no specific drugs or vaccines are available.

---

CoVs infect humans and other animal species, causing a variety of highly prevalent and severe diseases, including Severe Acute Respiratory Syndrome (SARS) and Middle East Respiratory Syndrome (MERS)^7^. The COVID-19 virus genome is comprised of ∼30,000 nucleotides; its replicase gene encodes two overlapping polyproteins, pp1a and pp1ab, required for viral replication and transcription^3,4^. The functional polypeptides are released from the polyproteins by extensive proteolytic processing, predominantly by a 33.8-kDa main protease (M^pro^), also referred to as the 3C-like protease. M^pro^ digests the polyprotein at no less than 11 conserved sites, starting with the autolytic cleavage of this enzyme itself from pp1a and pp1ab^8^. The functional importance of M^pro^ in the viral life cycle, together with the absence of closely related homologues in humans, identify the M^pro^ as an attractive target for antiviral drug design^9^.

To facilitate the rapid discovery of antiviral compounds with clinical potential, we developed a strategy combining structure-assisted drug design, virtual drug screening and high-throughput screening to repurpose existing drugs to target COVID-19 virus M^pro^.

## Establishing a high-throughput activity assay

Recombinant COVID-19 virus M^pro^ with native N and C termini was expressed in *Escherichia coli* and subsequently purified (Extended Data Fig. 1a, b). The molecular weight of COVID-19 virus M^pro^ as determined by mass spectroscopy is 33797.0 Da, consistent with its theoretical molecular weight 33796.8 Da. In order to characterize its enzymatic activity and to carry out high-throughput screening of inhibitors, we developed a fluorescence resonance energy transfer (FRET) assay. To do this, a fluorescently labeled substrate, MCA-AVLQ↓SGFR-Lys(Dnp)-Lys-NH_2_, derived from the N-terminal auto-cleavage sequence of the viral protease was designed and synthesized for time-dependent kinetic analysis (Extended Data Fig. 1e). The catalytic efficiency (*k_cat_/K_m_*) for COVID-19 virus M^pro^ was measured to be 28,500 M^−1^s^−1^ which is slightly higher than that for SARS-CoV M^pro^ (*k_cat_*/*K_m_*=26,500 M^−1^s^−1^)^10^, but over 30-fold higher than that of human rhinovirus 3C protease (*k_cat_*/*K_m_*=920 M^−1^s^−1^)^11^.

## N3 is a potent irreversible inhibitor of COVID-19 virus M^pro^

In a previous study, we designed a Michael acceptor inhibitor N3 using computer-aided drug design (CADD) (Extended Data Fig. 1c), which can specifically inhibit multiple CoV M^pro^s, including those from SARS-CoV and MERS-CoV^12–15^. It also has displayed potent antiviral activity against infectious bronchitis virus in an animal model^13^. The CC_50_ of N3 is >133 μM (Extended Data Fig. 1f). Next, we constructed a homology model for COVID-19 virus M^pro^ and used molecular docking to see if N3 could target this new CoV M^pro^. A docking pose showed that it could fit inside the substrate-binding pocket. To assess the efficacy of N3 for COVID-19 virus M^pro^, kinetic analysis was performed. A progress curve showed that it is a time-dependent irreversible inhibitor of this enzyme. Further, the shape of this curve supports the mechanism of two-step irreversible inactivation. The inhibitor first associates with COVID-19 virus M^pro^ (EI) with a dissociation constant *K_i_*; then, a stable covalent bond is formed between N3 and M^pro^ (E–I). The evaluation of this time-dependent inhibition requires both the equilibrium-binding constant *K_i_* (designated as *k_2_/k_1_*) and the inactivation rate constant for covalent bond formation *k*_3_. However, N3 exhibits very potent inhibition of COVID-19 virus M^pro^, such that measurement of *K_i_* and *k_3_* proved not feasible (Extended Data Fig. 1d, e). When very rapid inactivation occurs, *k_obs_*/[I] was utilized to evaluate the inhibition as an approximation of the pseudo second-order rate constant (*k_3_*/*K_i_*)^12^. The value of *k_obs_*/[I] of N3 for COVID-19 virus M^pro^ was determined to be 11,300±880 M^−1^s^−1^, suggesting this Michael acceptor has potent inhibition.

## The crystal structure of COVID-19 virus M^pro^ in complex with N3

In order to elucidate the inhibitory mechanism of this compound, we determined the crystal structure of COVID-19 virus M^pro^ in complex with N3 to 2.1-Å resolution. The asymmetric unit contains only one polypeptide (Extended Data Table 1). However, two of these associate to form a dimer by a crystallographic 2-fold symmetry axis (the two molecules are designated protomer A and B) (Fig. 1b). All residues (residues 1–306) are visible in electron density maps. Each protomer is composed of three domains (Fig. 1a). Domains I (residues 8–101) and II (residues 102–184) have an antiparallel β-barrel structure. Domain III (residues 201–303) contains five α-helices arranged into a largely antiparallel globular cluster, and is connected to domain II by means of a long loop region (residues 185–200). COVID-19 virus M^pro^ has a Cys–His catalytic dyad, and the substrate-binding site is located in a cleft between Domain I and II. These features are similar to those of other M^pro^s reported previously^5,6,13–15^. The electron density map shows that N3 binds in the substrate-binding pocket in an extended conformation (Fig. 1c, Extended Data Fig. 2), with the inhibitor backbone atoms forming an antiparallel sheet with residues 164–168 of the long strand_155-168_ on one side, and with residues 189–191 of the loop linking domains II and III.

**Fig. 1.**
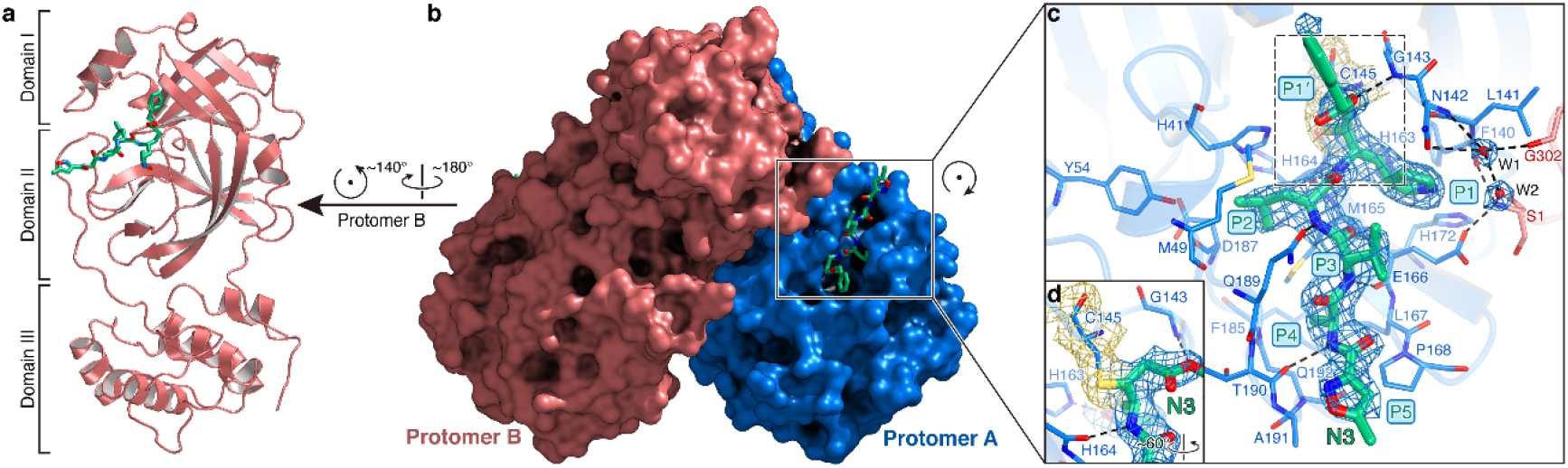
The crystal structure of COVID-19 virus M^pro^ in complex with N3. **a,** Cartoon representation of one protomer of the dimeric M^pro^-inhibitor complex. **b,** Surface representation of the homodimer of M^pro^. Protomer A is in blue, protomer B is in salmon, N3 is presented as green sticks. **c,** A zoomed view of the substrate-binding pocket. The key residues forming the binding pocket are shown in sticks, the two waters, assigned as W1 and W2, are shown as red spheres. P1, P1’, P2, P3, P4 and P5 sites of N3 are indicated. Hydrogen bonds that help to lock the inhibitor are shown in black dashed lines. The 2*F_o_-F_c_* density map contoured at 1.2 σ is shown around N3 molecule (blue mesh), C145-A (yellow mesh), and the two waters (blue mesh). **d,** The C-S covalent bond.

Here we detail the specific interactions of N3 with M^pro^ (Fig. 1c, d). The electron density shows that the Sγ atom of C145-A forms a covalent bond (1.8-Å) with the Cβ of the vinyl group, confirming that the Michael addition has occurred. The S1 subsite has an absolute requirement for Gln at the P1 position. The side chains of F140-A, N142-A, E166-A, H163-A, H172-A, S1-B (from protomer B), and main chains of F140-A and L141-A are involved in S1 subsite formation, which also includes two ordered water molecules (named W1 and W2). The lactam at P1 inserts into the S1 subsite and forms a hydrogen bond with H163-A. The side chain of Leu at P2 site inserts deeply into the hydrophobic S2 subsite, which consists of the side chains of H41-A, M49-A, Y54-A, M165-A, and the alkyl portion of the side chain of D187-A. The side chain of Val at P3 is solvent-exposed, indicating that this site can tolerate a wide range of functional groups. The side chain of Ala at P4 side is surrounded by the side chains of M165-A, L167-A, F185-A, Q192-A and the main chain of Q189-A, all of which form a small hydrophobic pocket. P5 makes van der Waals contacts with P168-A and the backbone of residues 190–191. The bulky benzyl group extends into the S1’ site, forming van der Waals interactions with T24-A and T25-A. In addition, N3 forms multiple hydrogen bonds with the main chain of the residues in the substrate-binding pocket, which also helps lock the inhibitor inside the substrate-binding pocket.

An overlay of the structures of COVID-19 virus M^pro^-N3 and SARS-CoV M^pro^-N1^12^ shows that N3 and N1 bind to M^pro^s in a similar mode (Fig. 2a, Extended Data Fig. 3). The major difference lies in the P1’ site. Compared with the benzyl ester portion of N3 in the COVID-19 virus M^pro^ structure, the ethyl ester portion in N1 adopts a slightly different conformation. This may be attributed to an ordered water (W1) in SARS-CoV M^pro^-N1 structure, which makes a long-distance hydrogen bond to the carboxylate oxygen of the ester and also forms two hydrogen bonds from the backbone NH of G143 and the side chain of N142. In our previous study, we proposed that all the CoV M^pro^s share a highly conserved substrate-recognition pocket, which could serve as a drug target for the design of broad-spectrum inhibitors^12^. The recent discovery of new CoVs and accumulation of structural data for CoV M^pro^s from various species provided the opportunity to further examine this hypothesis. Superposition of the 12 crystal structures of M^pro^s^12–21^ have shown that the most variable regions were the helical domain III and surface loops, but the substrate-binding pockets located in a cleft between domains I and II are still highly conserved among all CoV M^pro^s, suggesting the antiviral inhibitors targeting this site should have wide-spectrum anti-CoV activity (Fig. 2b, c).

**Fig. 2.**
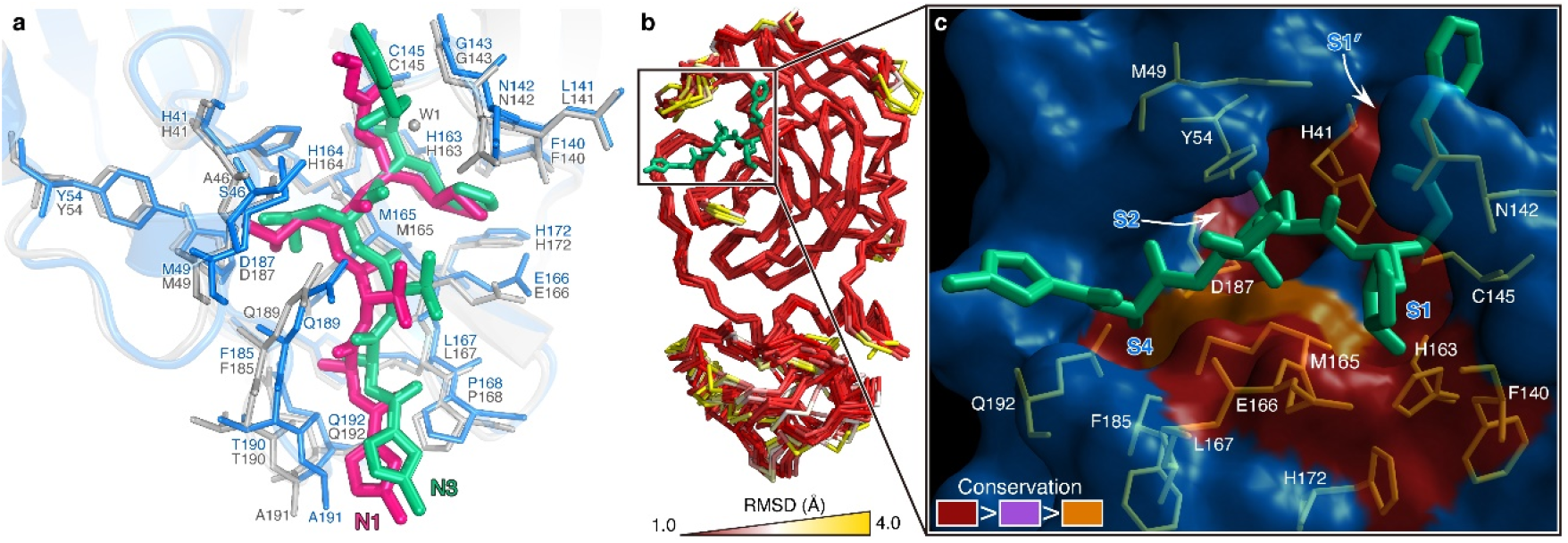
The substrate-binding pockets of CoV M^pro^s across different species. **a,** Comparison of inhibitor binding mode between the structures of COVID-19 virus M^pro^-N3 and SARS-CoV M^pro^-N1. COVID-19 virus M^pro^ is shown in marine cartoon; SARS-CoV M^pro^ in grey; N3 in green sticks; N1 in hot pink. **b,** Superposition of crystal structures of M^pro^s (Cα 1-300) from 12 CoVs, including COVID-19 virus, SARS-CoV, MERS-CoV, HCoV-HKU1, BtCoV-HKU4, MHV-A59, PEDV, FIPV, TGEV, HCoV-NL63, HCoV-229E and IBV The color spectrum represents the root-mean-square deviation (RMSD) of the aligned Cα atoms. **c,** Surface presentation of conserved substrate-binding pockets of 12 CoV M^pro^s. Red: residues are entirely identical among all 12 M^pro^s; violet: conserved substitution in one CoV M^pro^; orange: conserved substitution in more than one CoV M^pro^s. S1, S2, S4, and S1’ subsites are indicated.

## Virtual screening

The structure of COVID-19 virus M^pro^ in complex with N3 provides a model for identifying lead inhibitors to target COVID-19 virus M^pro^ through *in silico* screening. To achieve this, an in-house database of potential binding compounds was docked using Glide (v8.2). The results show that cinanserin fits snugly into the substrate-binding pocket, by interacting with H41 and E166 of M^pro^ by cation-π. Subsequently we determined this compound has an IC_50_ value of 125 μM for M^pro^. Moreover, cinanserin is a well-characterized serotonin antagonist, which underwent preliminary clinical testing in humans in the 1960s^22^ and has previously been shown to inhibit SARS-CoV M^pro23^. The CC_50_ of cinanserin is > 200 μM (Extended Data Fig. 4). Thus, it has potential for optimization as an anti-viral drug lead.

## High-throughput screening

Next, we used our FRET assay, to screen a library of ∼10,000 compounds consisting of approved drugs, clinical trial drug candidates and natural products. Primary hits included seven compounds that are either FDA-approved drugs or clinical trial/preclinical drug candidates. We then determined their IC_50_ values, which are in the range from 0.67 to 21.4 μM (Fig. 3). Amongst them, disulfiram and carmofur are FDA-approved drugs, whilst ebselen, shikonin, tideglusib, PX-12 and TDZD-8 are currently in clinical trials or preclinical studies. Ebselen has the strongest inhibition of M^pro^ activity with an IC_50_ of 0.67 μM. However, in a detergent-based assay^24^, TDZD-8 was found to be an aggregate-based inhibitor, which might non-specifically inhibit M^pro^ (Extended Data Fig. 5) and was therefore not considered for further investigation. Next, we set out to identify the potential covalent inhibitors among these compounds through tandem MS/MS analysis. The MS/MS data shows that ebselen, PX-12 and carmofur are all able to covalently bind to C145 of the catalytic dyad in COVID-19 virus M^pro^. However, while PX-12 and carmofur completely modified M^pro^, ebselen could only partially modify this viral cysteine protease (Extended Data Fig. 6). Since ebselen has even stronger inhibition than the others, there is a possibility that ebselen could also inhibit M^pro^ through non-covalent binding. It is likely that a portion of the hits identified by screening are covalently bonded to the catalytic cysteine of M^pro^ through their sulfhydryl groups. In general, such molecules are expected to be promiscuous binders and therefore, as they stand, may have limited potential as drug leads. Since our structural data is based on N3, we investigated if molecular docking could predict how disulfiram, tideglusib and shikonin bind to this protein. In all cases, reasonable docking poses were found, demonstrating that they could fit inside the substrate-binding pocket (Extended Data Fig. 7).

**Fig. 3.**
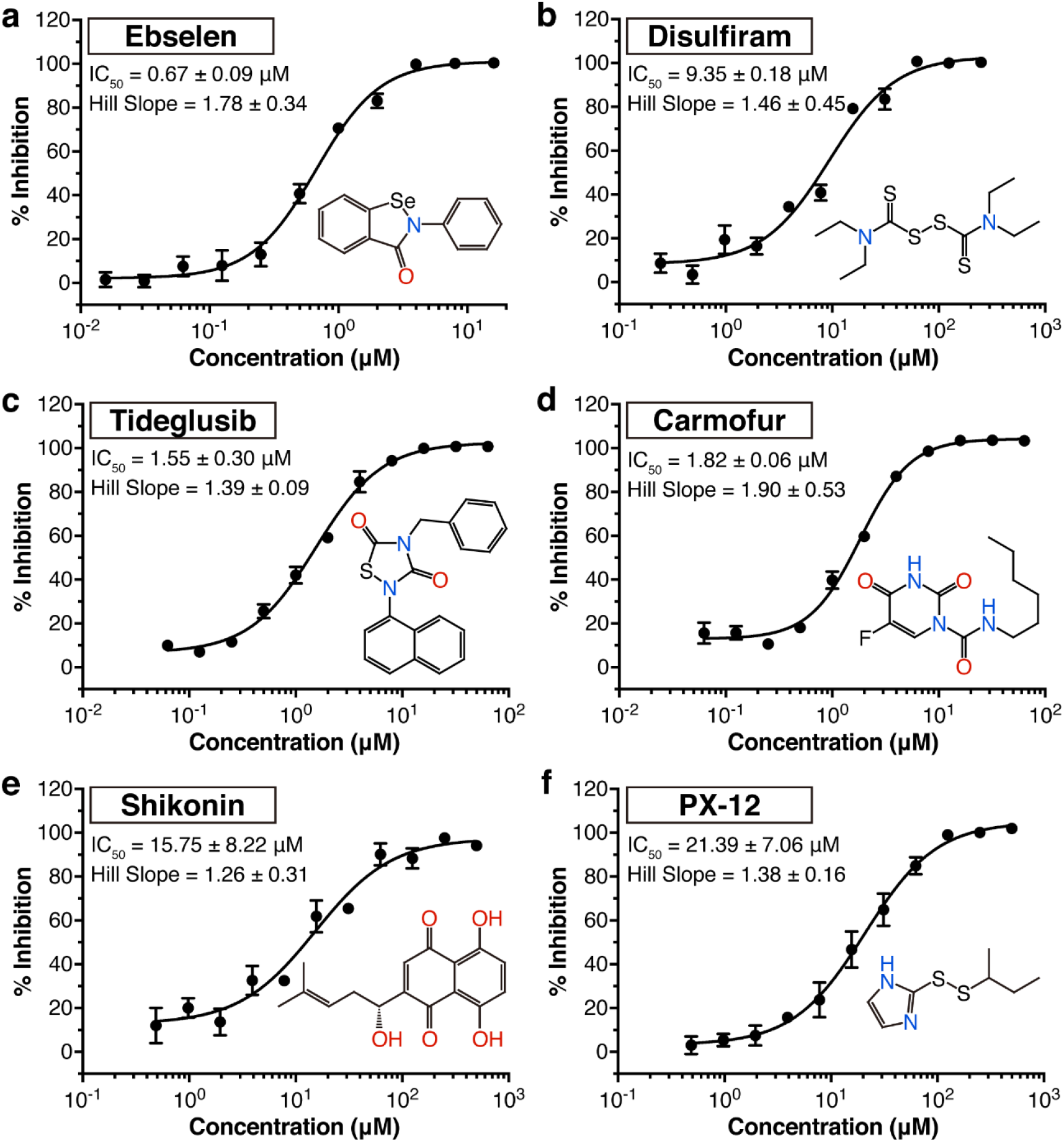
Drug leads inhibit the activity of COVID-19 virus M^pro^. **a-f,** The hydrolytic activity of COVID-19 virus M^pro^ was measured in the presence of varying concentrations of the drug candidates. Dose-response curves for half-maximum inhibitory concentration (IC_50_) values were determined by nonlinear regression. All data are shown as mean ± s.e.m., *n* = 3 biological replicates.

## Antiviral activity assay

To further substantiate the enzymatic inhibition results *in vitro*, we evaluated whether these compounds could prevent viral replication in cell-based assays. As shown in Fig. 4a, quantitative real-time RT-PCR (qRT-PCR) demonstrated that ebselen and N3 showed the strongest antiviral effects among them at a concentration of 10 μM treatment in COVID-19 virus infected Vero cells. A plaque-reduction assay (Extended Data Fig. 8) was performed to further assess the efficacy of these two compounds in protecting cells. Ebselen and N3 displayed inhibition against COVID-19 virus with individual EC50 values of 4.67 μM and 16.77 μM, respectively (Fig. 4b, c). The dose-response curves suggest that both of them could be able to penetrate cellular membrane to access their targets. Ebselen is an organoselenium compound with anti-inflammatory, anti-oxidant and cytoprotective properties. This compound has been investigated for the treatment of multiple diseases, such as bipolar disorders^25^ and hearing loss^26,27^. Ebselen has extremely low cytotoxicity (LD50 in rats > 4,600 mg/kg, per os)^28^ and its safety in humans has been evaluated in a number of clinical trials^26,27,29^. These data strongly suggest the clinical potential of ebselen for CoV treatment. It is also interesting that cinanserin displayed moderate inhibition against COVID-19 virus with an EC50 value of 20.61 μM from qRT-PCR analysis (Extended Data Fig. 4), which is superior to that in the enzymatic inhibition assay, suggesting that cinanserin might have multi-drug targets in preventing viral infection. In further studies, selection and characterization of drug-resistant mutants will help clarify the mode of cinanserin’s action.

**Fig. 4.**
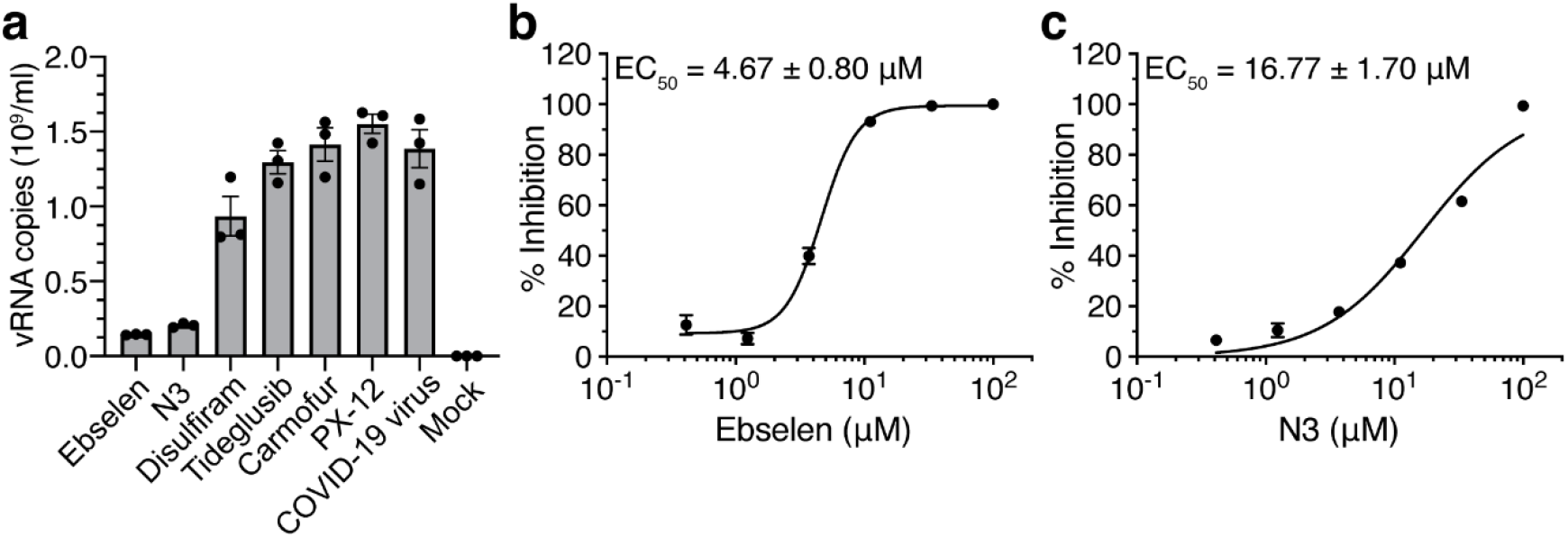
Antiviral activities of the drug leads against COVID-19 virus. **a,** The quantification of absolute viral RNA copies (per ml) in the supernatant at 72 h post infection (p.i.) determined by qRT-PCR analysis. Data are mean ± s.e.m., *n* = 3 biological replicates. **b, c,** Dose-response curves for ebselen and N3 in the plaque-reduction assay, respectively; all data are shown as mean ± s.e.m., *n* = 4 biological replicates.

## Discussion

Our crystal structural and docking data have shown that the drug leads identified can bind to the substrate-binding pocket of COVID-19 virus M^pro^, which is highly conserved among all CoV M^pro^s. This strongly supports our hypothesis that development of a single antiviral agent targeting M^pro^ or in combination with other potential therapies could provide an effective first line of defense against all CoV-associated diseases.

In the last twenty years, new infectious agents have emerged to cause epidemics, such as SARS and MERS^7^. The timely development of effective antiviral agents for clinical use is extremely challenging because conventional drug development approaches normally take years of investigations and cost billions of dollars. The repurposing of approved pharmaceutical drugs and drug candidates provides an alternative approach to rapidly identify potential drug leads to manage rapidly emerging viral infections. Cell-based phenotypic screening has proven to be valuable^30^, but the complexity of this approach is not readily compatible with high-throughput pipelines, and it cannot identify the molecular target or mechanism of action^31^. In this study, the convergence of structure-based *ab initio* drug design, virtual screening and high-throughput screening proved to be an efficient strategy to find antiviral leads against COVID-19 virus. The methods presented here can greatly assist in the rapid discovery of drug leads with clinical potential in response to new emerging infectious diseases that currently lack specific drugs and vaccines.

## Methods

### Cloning, protein expression and purification of COVID-19 virus M^pro^

The full-length gene encoding COVID-19 virus M^pro^ (NC_045512) was optimized and synthesized for *Escherichia coli* expression (Genewiz, USA). Cloning strategy for producing authentic viral M^pro^ has been reported previously^10^. The expression plasmid was transformed into *Escherichia coli* BL21 (DE3) cells and then cultured in Luria Broth medium containing 100 μg/ml ampicillin at 37 °C. When the cells were grown to OD_600_ of 0.6-0.8, 0.5 mM IPTG was added to the cell culture to induce the expression at 16 °C. After 10 h, the cells were harvested by centrifugation at 3,000*g*. The cell pellets were resuspended in lysis buffer (20 mM Tris-HCl pH 8.0, 300 mM NaCl), lysed by high-pressure homogenization, and then centrifuged at 25,000*g* for 40 min. The supernatant was loaded onto Ni-NTA affinity column (Qiagen, Germany), and washed by the resuspension buffer containing 20 mM imidazole. The His tagged M^pro^ was eluted by cleavage buffer (50 mM Tris-HCl pH 7.0, 150 mM NaCl) including 300 mM imidazole. Human rhinovirus 3C protease was added to remove the C-terminal His tag. The M^pro^ was further purified by ion exchange chromatography and size exclusion chromatography. CoV M^pro^s exist as a mixture of monomers and dimers in solutions^32^. The purified M^pro^ was stored in 50 mM Tris-HCl pH 7.3, 1 mM EDTA.

### Crystallization, data collection and structure determination

COVID-19 virus M^pro^ was incubated with 10 mM N3 for 30 min and the complex (5 mg/ml) was crystallized by hanging drop vapor diffusion method at 20 °C. The best crystals were grown with well buffer containing 0.1 M MES pH 6.0, 2% polyethylene glycol (PEG) 6000, 3% DMSO, 1 mM DTT. The cryo-protectant solution contained 0.1 M MES pH 6.0, 30% PEG 400.

X-ray data were collected on beamline BL17U1 at Shanghai Synchrotron Radiation Facility (SSRF) at 100 K and at a wavelength of 1.07180 Å using an Eiger X 16M image plate detector. Data integration and scaling were performed using the program Xia2^33^. The structure was determined by molecular replacement (MR) with the Phaser module^34^ in CCP4^35^ using the SARS-CoV M^pro^ (PDB ID: 2H2Z) as a search template. The output model from MR was subsequently subjected to iterative cycles of manual model adjustment with Coot^36^ and refinement was finished with Phenix^37^. The inhibitor N3 was built according to the omit map. The phasing and refinement statistics are summarized in Extended Data Table 1. The *R_work_/R_free_* values are 0.202/0.235, respectively. 97.3% residues are in most favored regions of the Ramachandran plot, and no residues are found in disallowed regions. Coordinates and structure factors for COVID-19 virus M^pro^ in complex with the inhibitor N3 have been deposited in Protein Data Bank (PDB) with accession number 6LU7. While this work was under review, we solved the complex structure at a higher resolution (1.7 Å). The relevant coordinates and structure factors have been deposited in PDB with accession number 7BQY.

### Enzymatic activity and inhibition assays

The enzyme activity assays have been described previously^10^. Briefly, the activity of COVID-19 virus M^pro^ was measured by a continuous kinetic assay, with the substrate MCA-AVLQSGFR-Lys(Dnp)-Lys-NH2 (GL Biochem, Shanghai), using wavelengths of 320 nm and 405 nm for excitation and emission, respectively. The assay started by immediately mixing 0.2 μM COVID-19 virus M^pro^ with different concentrations of substrate (2.5-100 μM). Fluorescence intensity was monitored with an EnVision multimode plate reader (Perkin Elmer, USA). Initial rates were obtained by fitting the linear portion of the curves to a straight line. The kinetic parameters *K_m_* and *k_cat_* were calculated from a double-reciprocal plot. As N3 is a mechanism-based irreversible inhibitor for COVID-19 virus M^pro^, *k_obs_*/[I] was used as an approximation of the pseudo second-order rate constant to evaluate the inhibition effect of the inhibitor N3^12^. In this case, the measurement was carried out with 0.2 μM of enzyme, 20 μM of substrate and inhibitor at 6 different concentrations (0-1 μM).

### Virtual screening

The virtual screening was performed using our in-house database via a workflow application of Glide (v8.2), Maestro (Schrödinger 2019-1a)^38^. All compounds in the database were considered to be at pH 7.4 ± 0.2 to estimate their protonation state using the program EpiK^39^. Their three dimensional conformations were generated by the ligPrep module of Maestro. The structure of COVID-19 virus M^pro^ (PDB ID: 6LU7) was used to generate receptor grid for docking simulations. The center of active site of the grid was determined according to the position of N3 in the structure. The flexibility of the receptor hydroxyl and thiol groups in side chains of C145, S46 and Y54 were considered. At the very beginning, a relatively fast but raw screening was performed by using the glide standard precision model, and the top 20% of compounds were kept. Finally, the candidate molecules were picked by analysing the predicted binding modes and their scores.

### High-throughput drug screen and IC_50_ measurement

Potential inhibitors against COVID-19 virus M^pro^ were screened by an enzymatic inhibition assay. When the different compounds were added into the enzymatic reaction mixture, the change of initial rates was calculated to evaluate their inhibitory effect. Five drug libraries, Approved Drug Library (Target Mol, USA), Clinic Compound Library (Target Mol, USA), FDA-approved Drug Library (Selleck, USA), Natural Product Library (Selleck, USA), and Anti-virus Drug Library (Shanghai Institute for Advanced Immunochemical Studies, SIAIS), which includes ∼10,000 compounds, were used. The preliminary screening reaction mixture included 0.2 μM protein, 20 μM substrate and 50 μM compounds. The compounds of interest were defined as those with a percentage of inhibition over 60% compared with the reaction in the absence of inhibitor. IC_50_ values of seven drug leads were measured using 0.2 μM protein, 20 μM substrate and 11 different inhibitor concentrations. In order to exclude inhibitors possibly acting as aggregators, detergent-based control was performed by adding 0.001% or 0.01% freshly made up Triton X-100 to the reaction at the same time^24^. All experimental data was analyzed using GraphPad Prism. All experiments were performed in triplicate.

### Molecular docking

To understand the binding interaction of these molecules with COVID-19 virus M^pro^, two different molecular docking methods, i.e., Glide (v8.2)^38^ and iFitDock^40^ were used to predict their binding poses. Then a 3D molecular similarity calculation method, SHAFTS^41^, was used for molecular alignment poses enumeration by matching the critical pharmacophore and volumetric overlay between the N3 molecule within the M^pro^ structure and the six drug candidates. However, the selenium atom of ebselen could not be treated by any of these above methods, so sulfur was used to replace it in the calculations. Then the obtained optimal superposition of these molecules was used to assess the reasonability of the predicted binding poses from the two docking methods, and only the binding orientations which were consistent among different methods were kept for constructing the initial complexes. Finally, these complexes were further optimized and re-scored by using MM-GBSA module^42^ of Schrödinger, and the residues within 5 Å around the ligand were refined.

### Antiviral and cytotoxicity assays for compounds from high-throughput screening

The *in vitro* antiviral efficacy of the drug candidates on Vero cells were determined by qRT-PCR. About 1×10^4^ Vero cells were seeded into a 96-well plate and incubated for 20-24 h at 37 °C. All the infection experiments were performed at biosafety level-3 (BSL-3). Cells were pre-treated with the drug candidates (10 μM) for 1 h; the COVID-19 virus (MOI of 0.01) was subsequently added to allow infection for 2 h. Then, the virus-drug mixture was removed and cells were further cultured with fresh drug-containing medium. At 72 h post infection (p.i.), viral RNA (vRNA) was extracted from the culture supernatant using QIAamp viral RNA mini kit (Qiagen, Germany) according to the manufacturer’s recommendation and detected by qRT-PCR assay using the COVID-19 virus-specific primers. Because shikonin showed cellular toxicity at the test concentration, its antiviral activity assay did not further proceed. Viral RNA copies per milliliter were determined using a synthetic RNA fragment to amplify the target region. The linearized plasmid containing S gene of COVID-19 virus was subjected to *in vitro* transcription. The resulting RNA transcripts were purified and then quantified using spectrophotometry on Nanodrop 2000 (Thermo Fisher Scientific, USA). The purified RNA was diluted 10-fold serially using RNase-free water and was detected using qRT-PCR. Threshold cycle (Ct) values for the known concentrations of the RNA were plotted against the log of the number of genome equivalent copies. The resultant standard curve was used to determine the number of genome equivalents of vRNA in the samples. The determination of the detection limit was based on the lowest level at which vRNA was detected and remained within the range of linearity of a standard curve (Ct value of 38). TaqMan primers for COVID-19 virus are 5’-TCCTGGTGATTCTTCTTCAGG-3’ and 5’-TCTGAGAGAGGGTCAAGTGC-3’ with COVID-19 virus probe 5’-FAM-AGCTGCAGCACCAGCTGTCCA-BHQ1-3’. The cytotoxicity of the tested drugs on Vero cell were determined by MTS cell proliferation assays (Promega, USA). 1×10^4^ cells were seeded into a 96-well plate and incubated for 20-24 h at 37 °C. After that, the medium was removed, and 100 μl of medium containing decreasing concentrations of antiviral compounds were added to the wells. After 4 days incubation at 37 °C, MTS assays were performed according to manufacturer’s protocols. All experiments were performed in triplicate. Vero cells were obtained from ATCC (American Type Culture Collection) with authentication service. All cell lines were tested negative for mycoplasma contamination. No commonly misidentified cell lines were used.

### Antiviral and cytotoxicity assays for cinanserin

For the antiviral assay, a clinical isolate COVID-19 virus^3^ was propagated in Vero E6 cells, and viral titer was determined as described previously^43^. All the infection experiments were performed at BSL-3. Pre-seeded Vero E6 cells (5×10^4^ cells/well) were pre-treated with the different concentrations of cinanserin for 1 h and the virus was subsequently added (MOI of 0.05) to allow infection for 2 h. Then, the virus-drug mixture was removed and cells were further cultured with fresh drug containing medium. At 24 h p.i., the cell supernatant was collected and vRNA in supernatant was subjected to qRT-PCR analysis. For cytotoxicity assays, Vero E6 cells were suspended in growth medium in 96-well plates. The next day, appropriate concentrations of cinanserin were added to the medium. After 24 h, the relative numbers of surviving cells were measured by CCK8 (Beyotime, China) assay in accordance with the manufacturer’s instructions. All experiments were performed in triplicate. Vero E6 cells were obtained from ATCC with authentication service. All cell lines were tested negative for mycoplasma contamination. No commonly misidentified cell lines were used.

### Plaque-reduction assays

1×10^5^ Vero E6 cells were seeded in a 24-well plate and treated with different doses of the inhibitors. All the infection experiments were performed at BSL-3. Inhibitors with different dilution concentrations were mixed with COVID-19 virus (100 PFU), 200 μl mixtures were inoculated onto monolayer Vero E6 cells for 1 h. After removing the supernatant, the plate was washed twice with DMEM medium, cells were incubated with 0.9% agarose containing appropriate concentrations of inhibitors. The overlay was discarded at 4 days p.i. and cells were fixed for 30 min in 4% polyoxymethylene and stained with crystal violet working solution. The plaque forming units were determined. All experiments were performed in four biological replicates.

### Intact protein analysis

2.5 μl of compounds (10 mM in DMSO) were added into 50 μl of COVID-19 virus M^pro^ (10 mg/ml). The mixtures were kept in room temperature for 30 min. Liquid chromatography-mass spectrometry (LC-MS) analyses were performed in positive-ion mode with a quadrupole-time-of-flight (QTOF) mass spectrometer (Agilent 6550, USA) coupled with a high-performance liquid chromatograph (HPLC, Agilent 1260, USA) for detecting the molecular weight of intact proteins. The samples were eluted from a Phenomenex Jupiter C4 300Å LC Column (2×150 mm, 5 μm) over a 15 min gradient from 5% to 100% acetonitrile containing 0.1% formic acid at a flow rate of 0.5 ml/min. The acquisition method in positive-ion mode with Dual Agilent Jet Stream electrospray voltage used a capillary temperature of 250 °C, a fragmentor of 175 V, a capillary voltage of 3000 V. Mass deconvolution was performed using Agilent MassHunter Qualitative Analysis B.06.00 software with BioConfirm Workflow.

### Tandem MS/MS analysis

The samples were precipitated and redissolved by 8 M urea, and then digested for 16 h at 25 °C by chymotrypsin at an enzyme-to-substrate ratio of 1:50 (w/w). The digested peptides were desalted and loaded onto a homemade 30 cm-long pulled-tip analytical column (ReproSil-Pur C18 AQ 1.9 μm particle size, Dr. Maisch GmbH, 75 μm ID× 360 μm OD) connected to an Easy-nLC1200 UHPLC (Thermo Fisher Scientific, USA) for mass spectrometry analysis. The elution gradient and mobile phase constitution used for peptide separation were as follows: 0-1 min, 4%-8% B; 1-96 min, 8-35% B; 96-104 min, 35-60% B; 105-120min, 60-100% B (mobile phase A: 0.1% formic acid in water; mobile phase B: 0.1% formic acid in 80% acetonitrile) at a flow rate of 300 nl /min. Peptides eluted from the LC column were directly electro-sprayed into the mass spectrometer with the application of a distal 1.8-kV spray voltage. Survey full-scan MS spectra (from m/z 300–1800) were acquired in the Orbitrap analyzer (Q Exactive, Thermo Fisher Scientific, USA) with resolution r = 70,000 at m/z 400. The top 20 MS/MS events were sequentially generated and selected from the full MS spectrum at a 30% normalized collision energy. The dynamic exclusion time was set at 10 seconds. One acquisition cycle includes one full-scan MS spectrum followed by top 20 MS/MS events, sequentially generated on the first to the twentieth most intense ions selected from the full MS spectrum at a 28% normalized collision energy. The acquired MS/MS data were analyzed UniProtKB *E.coli* database (database released on Nov. 11, 2016) containing nsp5 using Protein Discoverer 2.1. In order to accurately estimate peptide probabilities and false discovery rates (FDR), we used a decoy database containing the reversed sequences of all the proteins appended to the target database. FDR was set at 0.01. Mass tolerance for precursor ions was set at 20 ppm. Chymotrypsin was defined as cleavage enzyme and the maximal number of missed cleavage sites was set at 4. Protein N-terminus acetylation, methionine oxidation and compounds covalent bindings were set as variable modifications. The modified peptides were manually checked and labeled.

## Acknowledgments

We would like to thank Ying Lei and Juan Kong from High Throughput Platform, staff from Analytical Chemistry Platform at Shanghai Institute for Advanced Immunochemical Studies, for their technical support. We are grateful to National Centre for Protein Science Shanghai and The Molecular and Cell Biology Core Facility of the School of Life Science and Technology, ShanghaiTech University for use of their instrumentation and technical assistance. We thank Prof. Zhijie Liu, and Haixia Su for discussion. We also thank the staff from beamlines BL17U1, BL18U1 and BL19U1 at Shanghai Synchrotron Radiation Facility (China). This work was supported by grants from National Key R&D Program of China (grant No. 2017YFC0840300 to Z.R.), Project of International Cooperation and Exchanges NSFC (grant No. 81520108019 to Z.R.), Science and Technology Commission of Shanghai Municipality (grant No. 20431900200) and Department of Science and Technology of Guangxi Zhuang Autonomous Region (grant No. 2020AB40007).

## Author contributions

Z.R. and H.Y. conceived the project; Z.J., H.J., Z.R., and H.Y. designed the experiments; Z.J., X.D., Y.Duan., J.Y., T.Y., Xiaoce Liu and Xiuna Yang cloned, expressed, purified and crystallized proteins; Z.J., Y.Z., B.Z. and F.L. collected the diffraction data; B.Z. and Xiang Liu solved the crystal structure; Z.J., X.D., Y.Duan. and J.Y. performed enzymatic activity and inhibition assay, high-throughput drug screen and IC_50_ measurement; L.W. and F.B. performed virtual screening and molecular docking; Y.X., L.Z. and H.L. performed enzymatic inhibition, cell-based antiviral and cytotoxicity assay for cinanserin; Y.Deng. and X. Li performed qRT-PCR analysis and cytotoxicity assay of N3; M.L., R.J. and Xinglou Yang performed plaque-reduction assay; C.P. performed intact protein and tandem MS/MS analyses; Z.J., X.D., Y.X., Y.Deng., C.P., F.B., H.L., Xiang Liu, K.Y., L.G., W.X., G.X., C.Q., Z.S., H.J., Z.R. and H.Y. analyzed and discussed the data; Z.J., X.D., F.B., Xiang Liu, L.G., G.X., C.Q., Z.S., H.J., Z.R. and H.Y wrote the manuscript.

## Data and materials availability

The PDB accession No. for the coordinates of COVID-19 virus M^pro^ in complex with N3 is 6LU7 (Deposited: 2020-01-26; Released: 2020-02-05).

---

**Extended Data Fig. 1.**
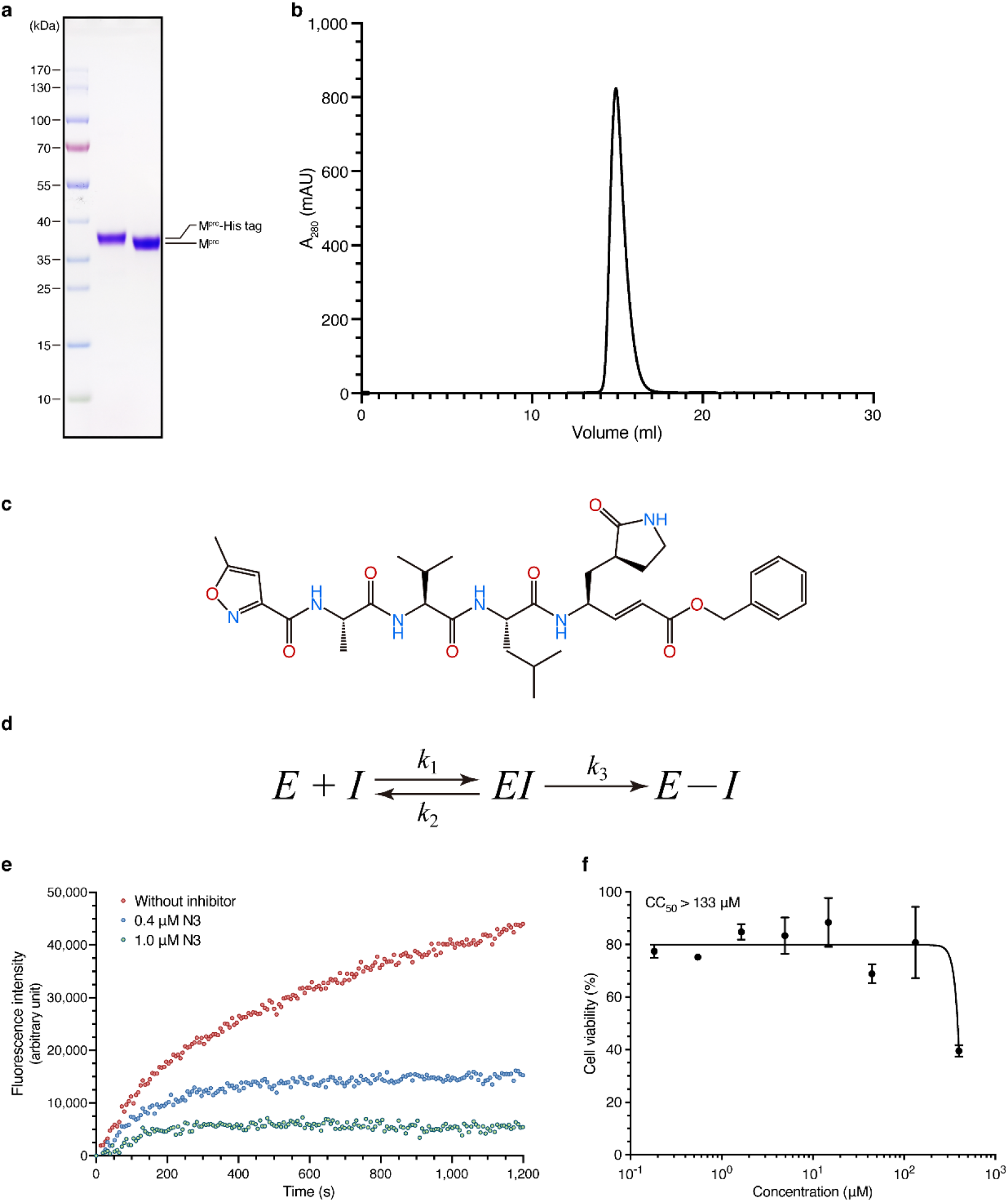
The purification of COVID-19 virus M^pro^ and the inhibitory assay of N3 compound. **a,** The SDS-PAGE gel of COVID-19 virus M^pro^. The first lane: marker; the second lane: M^pro^ before treating with rhinovirus 3C protease; third lane: M^pro^ after the cleavage of C-terminal His tag. For gel source data, see Supplementary Figure 1. **b,** Size-exclusion chromatography profile of M^pro^. **c,** The chemical structure of N3 inhibitor. **d,** Inhibition mechanism for N3. **e,** Typical inhibition curves for N3. **f,** Cytotoxicity assay of N3 on Vero cells, data are shown as mean ± s.e.m., *n* = 3 biological replicates. The data (**a, b, f**) are representative of three independent experiments with similar results.

**Extended Data Fig. 2.**
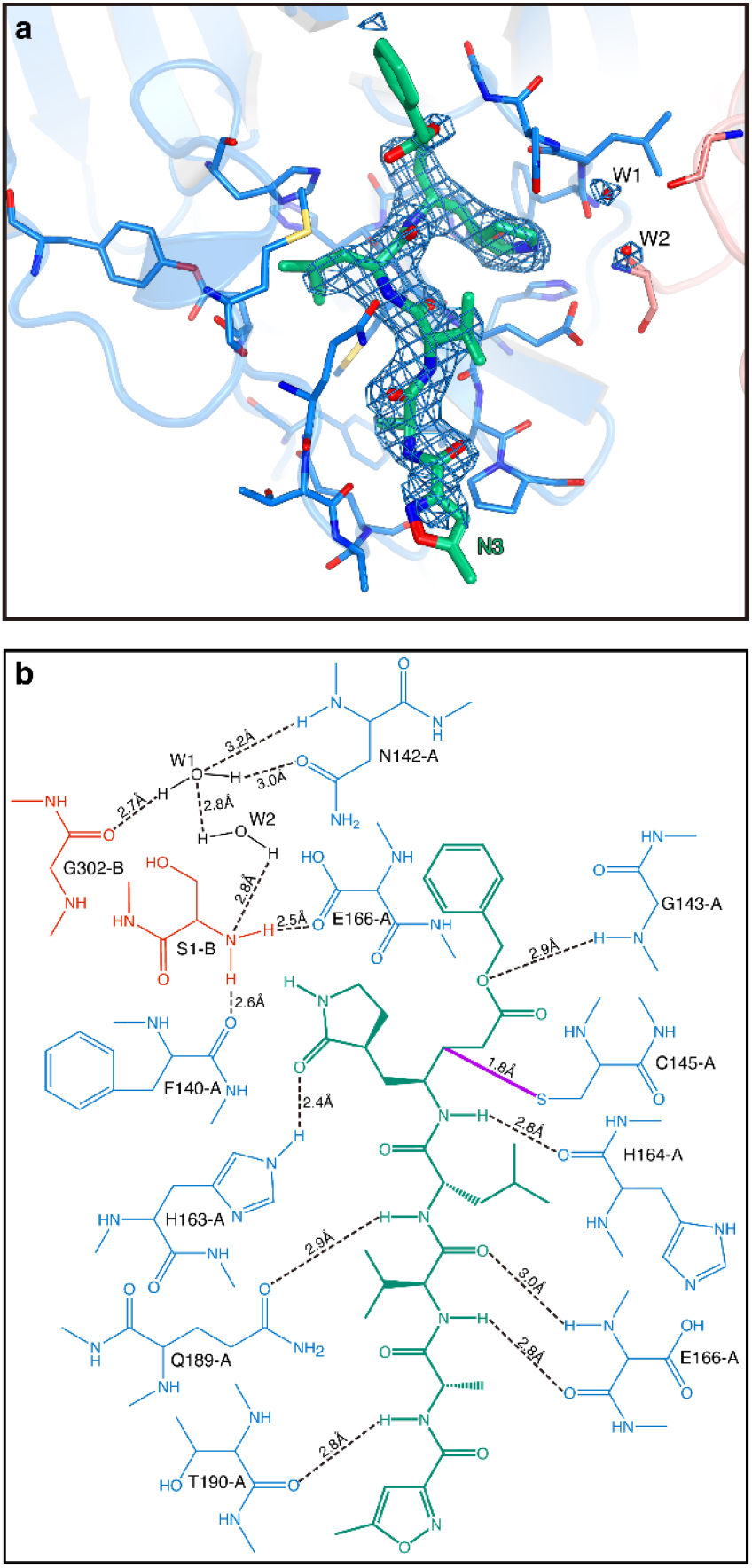
The interactions between COVID-19 virus M^pro^ and N3. **a,** The *F_o_-F_c_* omit map (contour level = 3 σ, shown as the blue mesh). **b,** Detailed view of the interactions between the inhibitor N3 and COVID-19 virus M^pro^. M^pro^ residues are shown in blue (Protomer A) and salmon (Protomer B); N3 is in green, water is in black. The hydrogen bonds are shown as black dashed lines. The covalent bond between N3 and C145-A is in purple.

**Extended Data Fig. 3.**
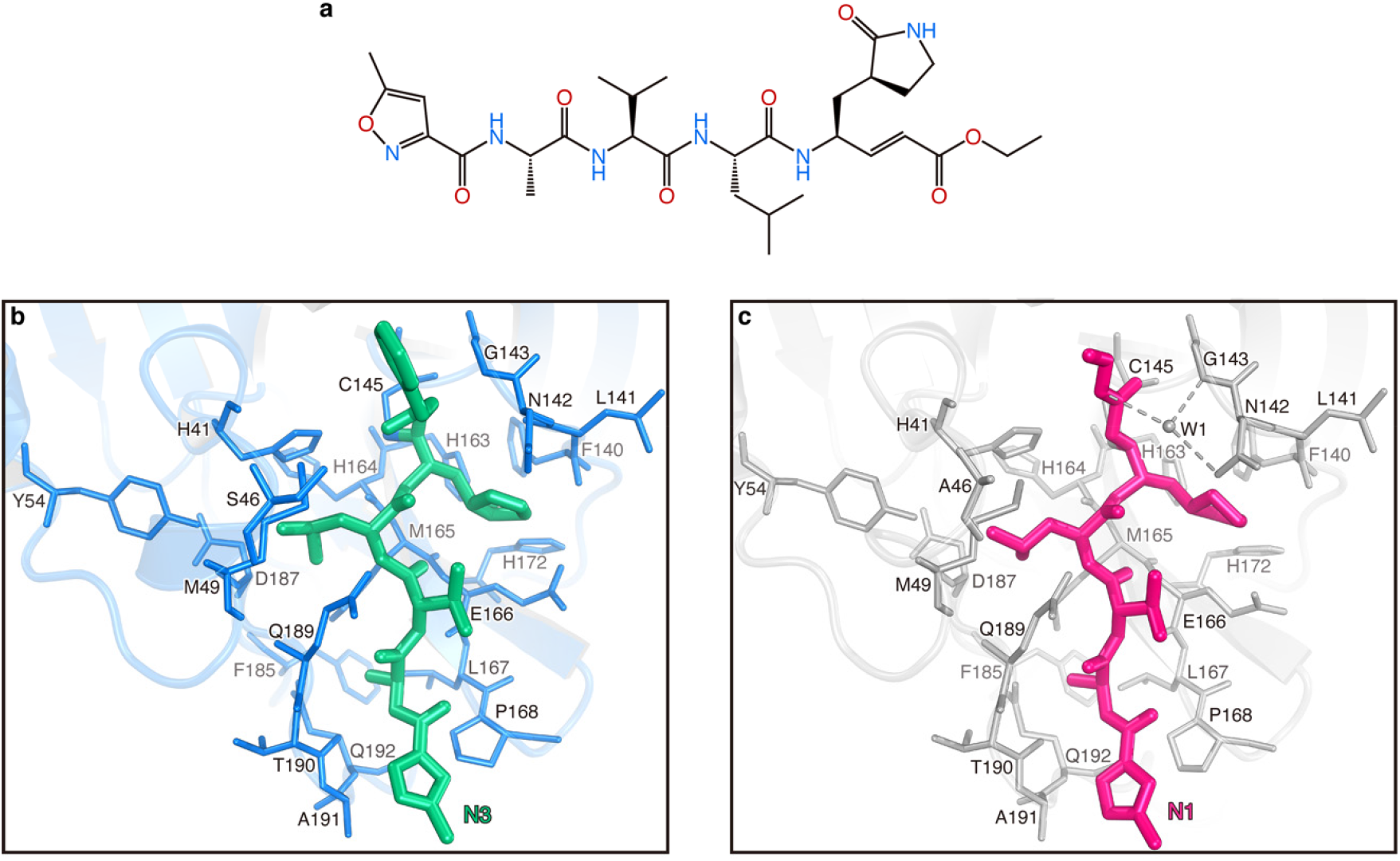
Comparison of the binding modes between COVID-19 virus M^pro^-N3 and SARS-CoV M^pro^-N1. **a,** The chemical structure of N1 inhibitor. **b,** The binding mode of COVID-19 virus M^pro^ (blue sticks) with N3 (green sticks). **c,** The binding mode of SARS-CoV M^pro^ (grey sticks) with N1 (pink sticks). The hydrogen bonds formed by water (W1) are indicated by the dashed lines.

**Extended Data Fig. 4.**
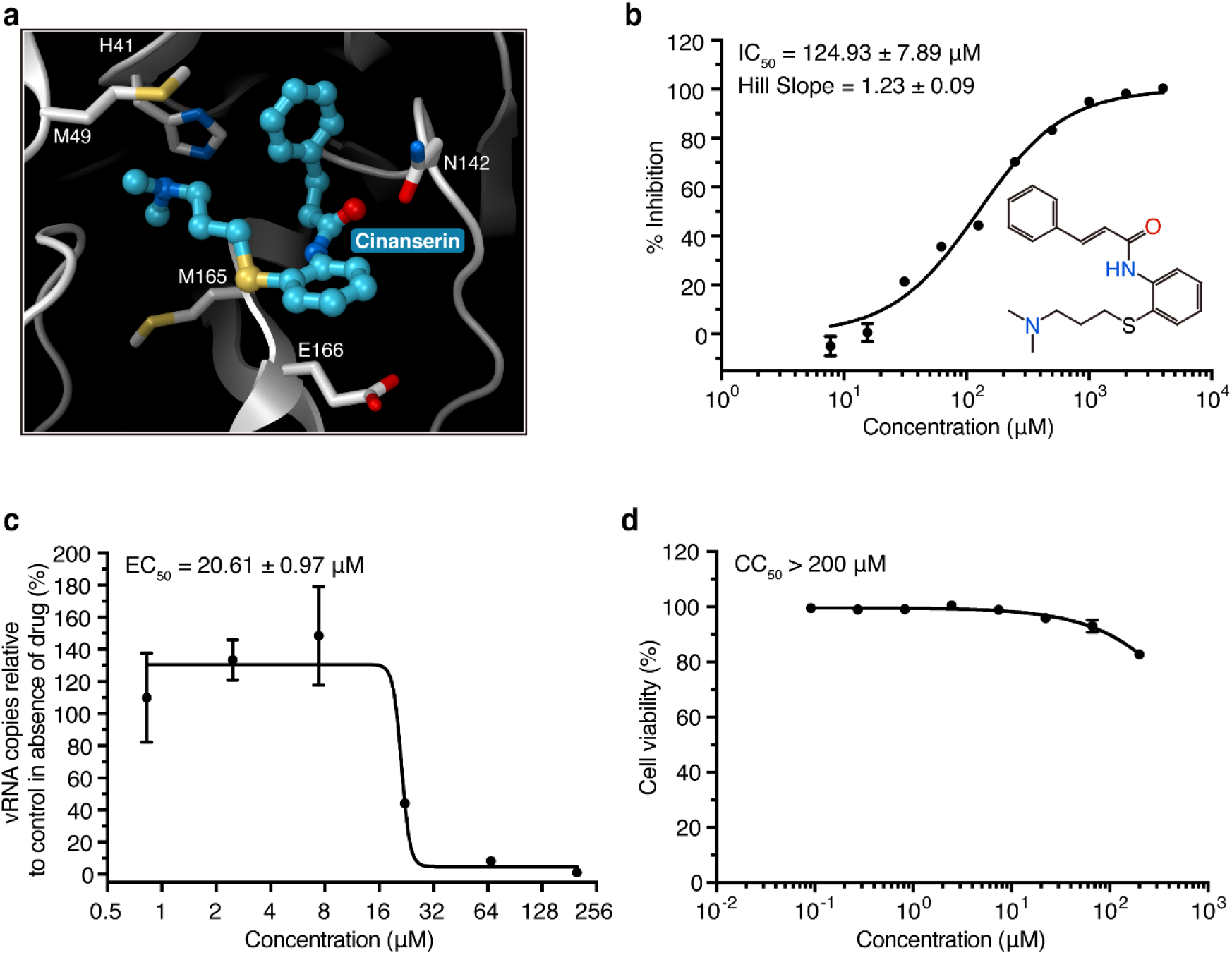
Cinanserin is an inhibitor for COVID-19 virus M^pro^. **a,** The docking result of cinanserin. The structure of COVID-19 virus M^pro^ is shown as a white cartoon, cinanserin is shown as cyan balls and sticks, residues predicted to be interacting with cinanserin are shown as sticks. **b,** Inhibitory activity of cinanserin on M^pro^. **c,** Antiviral activity of cinanserin determined by qRT-PCR. **d,** Cytotoxicity assay of cinanserin on Vero E6 cells. All data are shown as mean ± s.e.m., *n* = 3 biological replicates.

**Extended Data Fig. 5.**
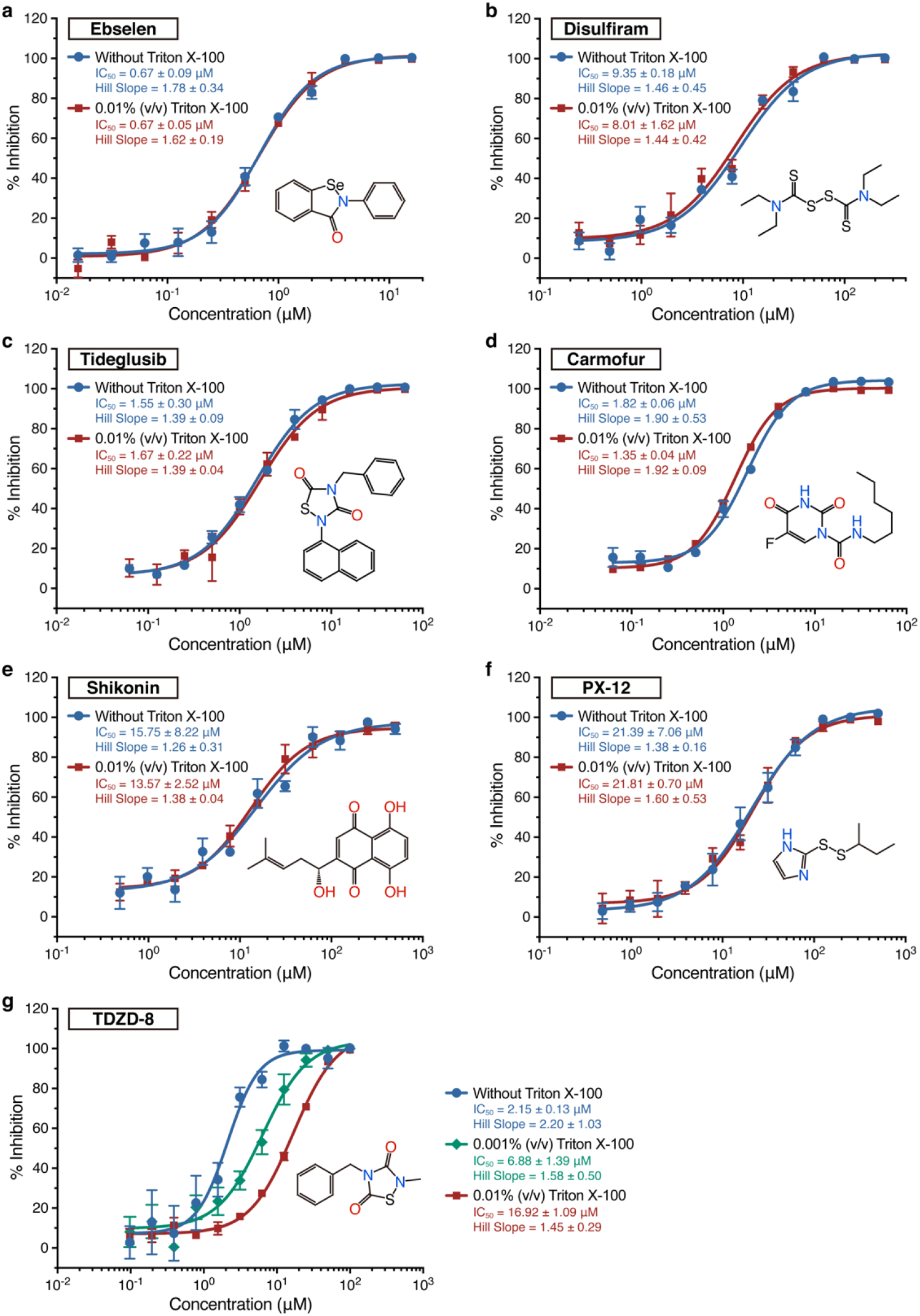
The detergent-based assay for drug leads. **a-f,** The IC_50_ values determined by in the presence or absence of 0.01% Triton X-100, which showed that detergent did not affect the results. **g,** Different concentrations of Triton X-100 notably affected IC_50_ curves for TDZD-8. All data are shown as mean ± s.e.m., *n* = 3 biological replicates.

**Extended Data Fig. 6.**
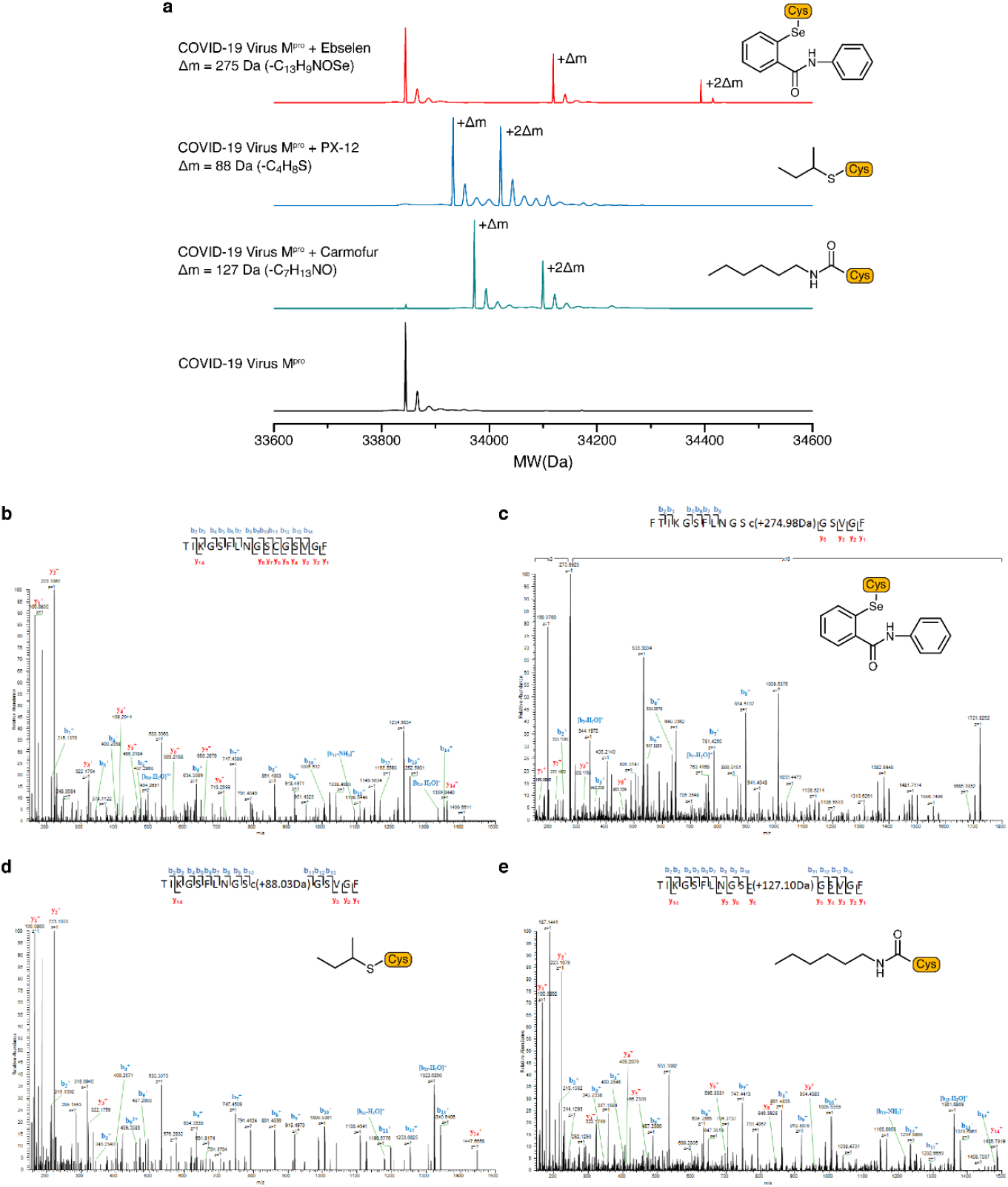
Tandem MS/MS analysis reveals that ebselen, PX-12 and carmofur are able to covalently bind to C145 of COVID-19 virus M^pro^. **a,** Molecular weight of apo COVID-19 virus M^pro^ and compounds treated M^pro^. The mass shifts (Δm) of the proteins indicate that more than one molecular of the compounds can be covalently bonded to one molecular of M^pro^. **b-e,** A higher-energy collisional dissociation (HCD) MS/MS spectrum recorded on the [M+H]^2+^ ion **b,** at m/z 787.3852 of the M^pro^ unmodified peptide TIKGSFLNGSCGSVGF, **c,** at m/z 998.4152 of the M^pro^ modified peptide FTIKGSFLNGSCGSVGF harboring a modification (-C_13_H_9_NOSe) induced by ebselen on C145, **d,** at m/z 831.4080 of the M^pro^ modified peptide TIKGSFLNGSCGSVGF harboring a modification(-C_4_H_8_S) induced by PX-12 on C145, **e,** at m/z 850.9414 of the M^pro^ modified peptide TIKGSFLNGSCGSVGF harboring a modification(-C_7_H_13_NO) induced by carmofur on C145. Predicted b- and y-type ions (not including all) are listed above and below the peptide sequence, respectively. The experiment was performed once.

**Extended Data Fig. 7.**
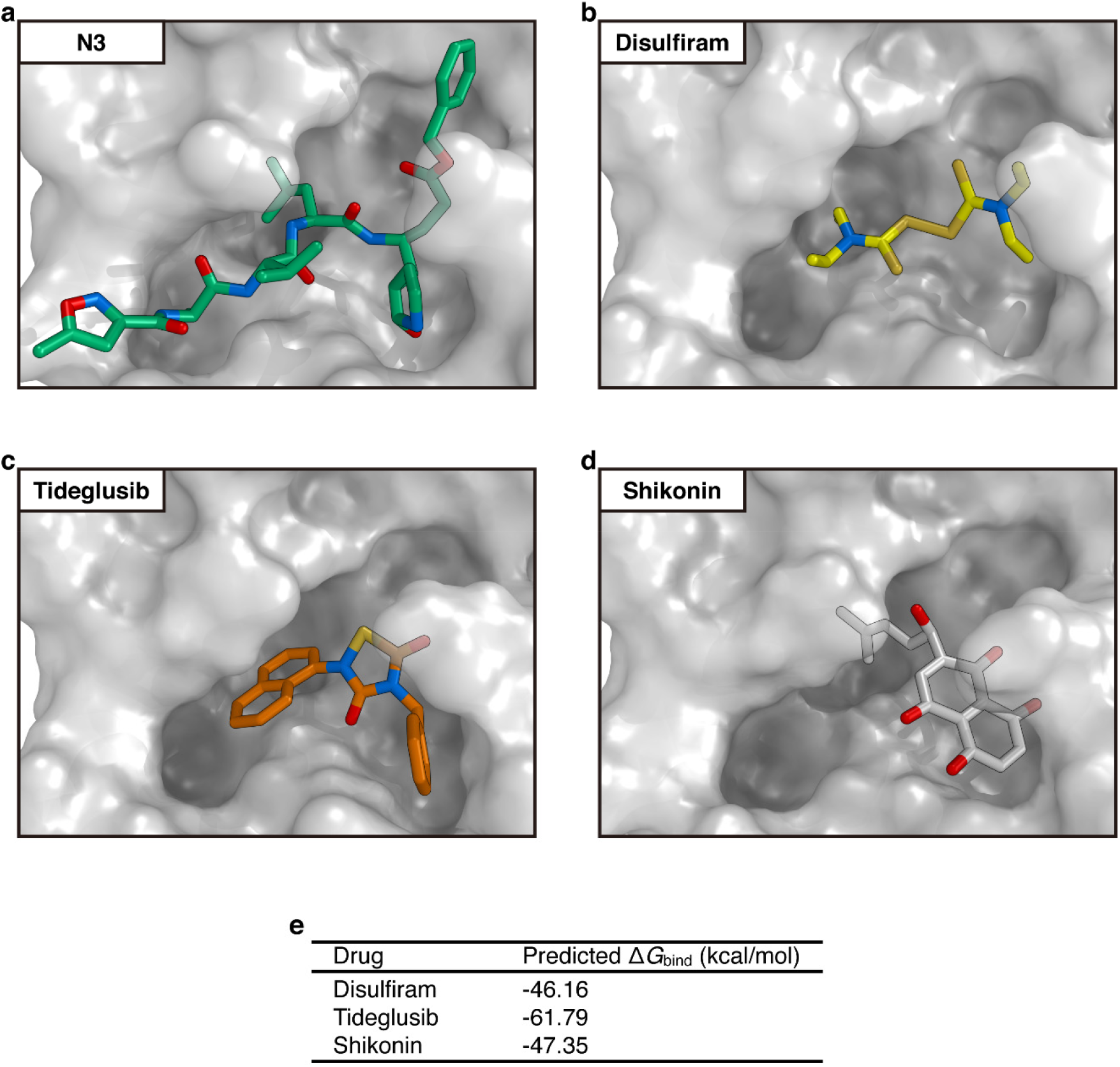
Docking Poses of different COVID-19 virus M^pro^ inhibitors. **a,** The crystal structure of COVID-19 virus M^pro^-N3 complex. **b-d,** The docking results of three drug leads. M^pro^ is shown as grey background, inhibitors are in different colors. The inhibitors identified through the high-throughput screening are likely to occupy the same pocket as N3. **e,** Predicted binding affinities for the drug leads to COVID-19 virus M^pro^ by using MM-GBSA module integrated in Schrödinger.

**Extended Data Fig. 8.**
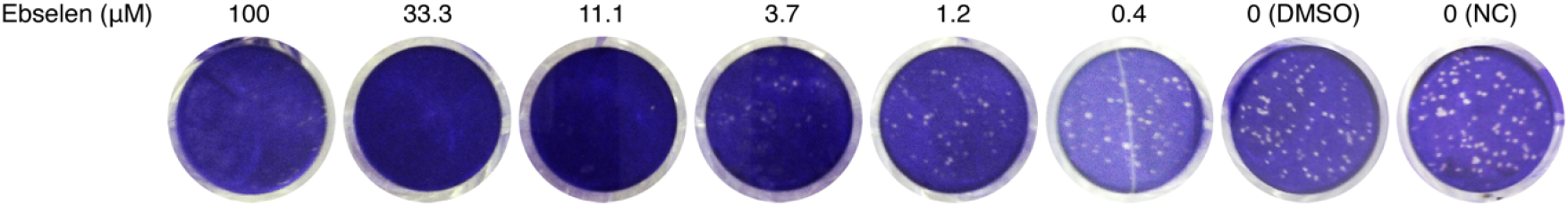
Images for the plaque-reduction assay (ebselen). As the concentration of the inhibitor (ebselen) increases, there is a significant reduction in the numbers of the plaques by comparison with NC (negative control) and DMSO. Results are shown as representative of four biological replicates. For image source data, see Supplementary Figure 2.

**Extended Data Table 1.**
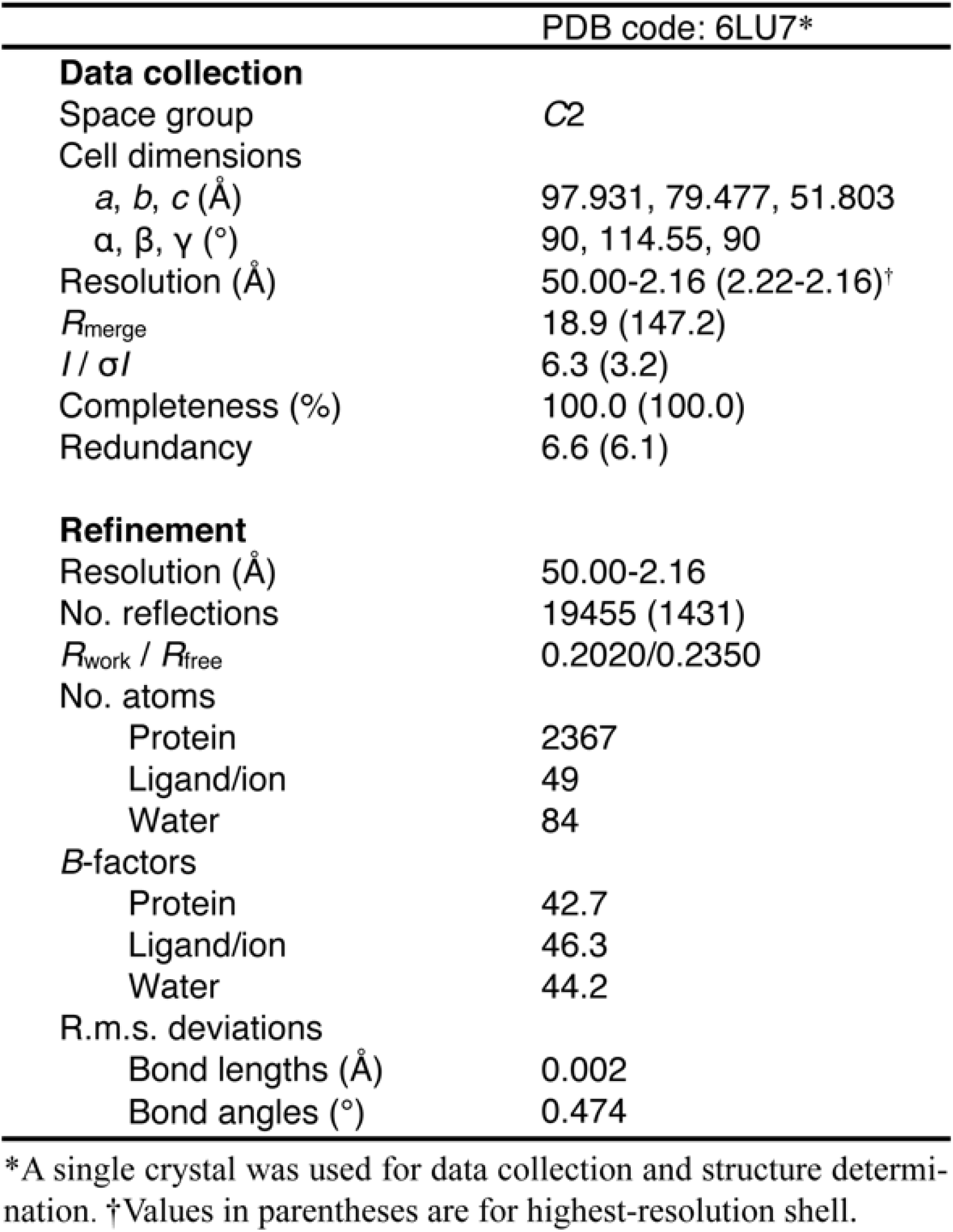
Data collection and refinement statistics.

